# Aryl hydrocarbon receptor (AHR) and IL-13 signaling crosstalk in human keratinocytes and atopic dermatitis

**DOI:** 10.1101/2023.06.17.545291

**Authors:** Steven P. Proper, Alexander T. Dwyer, Andrews Appiagyei, Jennifer M. Felton, Netali Ben-Baruch Morgenstern, Justin M. Marlman, Michael Kotliar, Artem Barski, Ty D. Troutman, Marc E. Rothenberg, Tesfaye B. Mersha, Nurit P. Azouz

## Abstract

Atopic dermatitis (AD) is an allergic skin disease mediated by skin barrier impairment and IL-13-driven immune response. Activation of the aryl hydrocarbon receptor (AHR) has shown promise in early clinical trials for AD, however, the mechanism by which AHR mediates this function is unknown. Herein, AHR signaling is shown to be dysregulated in AD patients using publicly available gene expression data from biopsies of patients with AD compared with controls. AHR target genes, *CYP1A1*, *CYP1A2* and *NFE2L2* were decreased in lesional skin compared with healthy control skin (p=0.001, p<1.0*10^-10^ and p=6.1*10^-6^ respectively). Single cell RNA sequencing studies demonstrated increased expression of *AHR* (p<1.0*10^-4^ and p=0.049) and decreased expression of *CYP1A1* in lesional AD keratinocytes compared with healthy control keratinocytes (p=5.0*10^-4^ and p=0.21). AHR activation reversed IL-13-dependent gene expression of several key genes in AD pathogenesis, most notably the eosinophil chemoattractant, *CCL26* (Eotaxin-3). There was substantial overlap between differentially expressed genes in keratinocytes of patients with atopic dermatitis and AHR-regulated genes. Mechanistically, there was evidence for direct transcriptional effects of AHR as its binding motifs were identified in the differentially expressed genes from lesional AD keratinocytes compared to control keratinocytes and AHR activation did not modify IL-13-dependent signal transducer and activator of transcription 6 (STAT6) translocation to the nucleus. Together these data imply that AHR modulates IL-13 downstream signaling in keratinocytes through genome-wide direct transcriptional regulatory effects.

## Introduction

Atopic dermatitis (AD) is a chronic, relapsing inflammatory skin disorder with a hallmark of impaired skin barrier function affecting up to 11% of children and 7% of adults in the US and up to 20% of people worldwide. AD is considered to be a major risk factor for the development of other allergic diseases^1, 2^. A combination of environmental and genetic factors contribute to the local and systemic inflammation that drive AD. Evidence is mounting for the importance of AHR signaling in keratinocytes for promoting epidermal differentiation and preventing AD progression. Tapinarof, a naturally occurring (now fully synthetic) AHR agonist, has shown promise in treating AD through Phase 2 clinical trials (and Phase 3 clinical trials are underway)^3–5^. While AHR activation improves AD, it is still unclear whether these changes are driven largely by keratinocytes or other cell types. In addition, the mechanism by which AHR activation interferes with Type 2 inflammation seen in AD is not fully understood. Originally defined due to its role in mediating the toxicity of xenobiotics such as 2,3,7,8-tetrachlorodibenzo-p-dioxin (TCDD), AHR reacts to multiple skin-relevant ligands to promoting epithelial differentiation and regulating immune responses^6^. AHR endogenous ligands include 6-formyl-indolo[3,2-b]carbazole (FICZ), a product of tryptophan and ultraviolet radiation in the skin^7^, indoles from foods and microbiota^8^ and other tryptophan metabolites such as kynurenine^9^. These endogenous ligands of AHR are far more transient and subject to metabolic breakdown compared to TCDD and other toxicants such as polychlorinated biphenyls (PCBs) and benzo[A]pyrene. Further research continues to elucidate the cell-type-specific physiologic roles of the AHR ligands.

Keratinocytes are the primary functional cells of the epidermis, and form a physical barrier in addition to being active participants in directing innate immune responses. As the first line of defense from the outside environment, keratinocytes are uniquely positioned to have significant potential exposure to AHR ligands, whether through exposure to UV radiation, proximity to commensal skin microbiota, and direct exposure to particulate matter and products of combustion.

Herein, we present evidence that AHR signaling is dysregulated in keratinocytes from patients with AD using publicly available gene expression data. We found that AHR activation blocks IL-13-dependent expression of key genes in AD pathogenesis, including CCL26 (Eotaxin-3), a crucial cytokine regulated by STAT6. We showed substantial overlap between the AHR-regulated transcriptome and the AD transcriptome in keratinocytes. Analysis of STAT6 did not reveal substantial changes in STAT6 expression, phosphorylation, or translocation to the nucleus by AHR. In contrast, we demonstrated that among open chromatin regions from keratinocytes in AD lesional skin, there is an enrichment of AHR binding motifs near the transcriptional start site of differentially regulated genes. These data implicate DNA binding as the primary mechanism by which AHR activation alters type 2 signaling in keratinocytes in AD, and highlight the need for further study to elucidate these molecular mechanisms as AHR activation is utilized to treat AD.

## Methods

### Cell Culture and Reagents

HaCaT cells were obtained from CLS (item no. 300493, Eppelheim, Germany), and validated via STR profiling by Labcorp (Burlington, North Carolina, USA). HaCaT cells were cultured in Gibco DMEM from Thermo Fischer (Cat. No. 10567014, Waltham, MA, USA), supplemented with 10% Fetal Bovine Serum (FBS) from R&D Biosystems (Cat. No. S11150, Minneapolis, MN, USA) and 1% Penicillin/Streptomycin from Gibco/Thermo Fisher Scientific (Cat. No. 15140-122, Waltham, MA, USA). Cells were plated onto Falcon brand cell culture plates from Corning (Corning, NY, USA) and kept at 37°C and 5% CO_2_. Passaging of cells carried out with 0.05% trypsin from Gibco/Thermo Fisher Scientific (Cat. No. 25300-054, Waltham, MA, USA) and 10X phosphate buffered saline from Gibco/Thermo Fisher Scientific (Cat. No. 14200-075, Waltham, MA, USA) that was diluted to 1X with purified water from NanoPure filtration system (Thomas Scientific, Swedesboro, NJ, USA) that was then autoclave-sterilized. HaCaT cells were grown to confluence and then continued for 3 days to ensure monolayer was achieved prior to dosing. Dimethylsulfoxide (DMSO) was obtained from Sigma-Aldrich (Cat. No. D2650, St. Louis, MO, USA) and used to dilute FICZ, Tapinarof and GNF351 prior to their final dilution in culture media. FICZ was obtained from Tocris/Bio-Techne (Cat. No. 5304, Minneapolis, MN, USA), put into 1 mM stock solution with DMSO kept at −20°C and protected from light prior to use in experiments, when it was further diluted into culture media at 1:1000 to final concentration of 1 µM. Tapinarof was obtained from MedChemExpress (Cat. No. HY-109044, Monmouth Junction, NJ, USA), put into 1 mM stock solution with DMSO and similarly diluted with culture media to final concentration of 1 µM. GNF351 was obtained from EMD Millipore (Cat. No. 182707-10MG, Burlington, MA, USA), put into 1 mM stock solution with DMSO and diluted into culture media for final concentration of 1 µM. Unless otherwise noted, chemical reagents were obtained from Sigma-Aldrich (St. Louis, MO, USA). After dosing, cells were washed with 1X ice-cold PBS prior to isolation of either protein or RNA as described below.

### mRNA extraction and quantitative RT-PCR

Total RNA was isolated from cells using TriPure Isolation Reagent from Sigma-Aldrich using the manufacturer’s instructions (Cat. No. 11667165001, St. Louis, MO, USA). RNA layer was further purified using the RNeasy Mini Kit by Qiagen (Cat. No. 2170004, Germantown, MD, USA) according to manufacturer instructions. cDNA was created from RNA using Protoscript First Strand cDNA Synthesis Kit from New England BioLabs (Cat. No. E63005, Ipswitch, MA, USA). qPCR was performed using an ABI QuantStudio 7 Flex (Thermo Fisher, Waltham, MA, USA) with PowerUp SYBR Green Master Mix from Thermo Fisher (Cat. No. A25742, Waltham, MA, USA) using the following primer sets: GAPDH (forward 5’-AGGTCGGAGTCAACGGATTT, reverse 5’- GACGGTGCCATGGAATTTGC), CYP1A1 (forward 5’-AGTGATTGGCAGGTCACGG, reverse 5’-GTCTCTTGTTGTGCTGTGGGG), CCL26 (forward 5’- TCCCAGCGGGCTGTGATATTC, reverse 5’- TCCAAGCGTCCTCGGATGAA).

### Protein Extraction and Western Blot

Nuclear and Cytosolic protein fractions from cell cultures were extracted with NE-PER Nuclear and Cytoplasmic Extraction Reagents from Thermo Fisher (Cat. No. 78835, Waltham, MA, USA), and protein from whole cell fraction as well as supernatants were extracted with M-PER buffer from Thermo Fisher Scientific (Cat. No. 78501, Waltham, MA, USA) with protease inhibitors from Roche/Sigma-Aldrich (Cat. No. 11836153001, St. Louis, MO, USA). 4X Bolt LDS Sample Loading buffer from Invitrogen/Thermo Fisher (Cat. No. B0008, Waltham, MA, USA) was added, and samples were sonicated at 10 kHz for two 10-second intervals with a 5 second break in between intervals (Fisher Scientific UltraSonic Processor). Samples were then heated to 95°C for 5 min, and cellular debris spun down at 12,000 g for 5 minutes before being subjected to electrophoresis on Bolt 4- 12% Bis-Tris gels from Invitrogen/Thermo Fisher (Cat. No. NW04127BOX, Waltham, MA, USA) at 200 V for 30 minutes, then transferred to SureLock Tandem Midi Pre-cut nitrocellulose membranes from Invitrogen/Thermo Fisher (Cat. No. 11836153001, Waltham, MA, USA) at 30 V for 1 hour, and then visualized using the Odyssey CLx system (LI-COR Biosciences). Membranes were blocked with Intercept TBS Bocking Buffer from Li-Cor (Cat. No. 927-60001, Lincoln, NE, USA) prior to incubation with primary antibodies. Primary antibodies were rabbit anti-AHR monoclonal IgG (Cell Signaling 83200, clone D5S6H, 1:2000) and rabbit anti-pSTAT6 (Tyr641) monoclonal IgG (Cell Signaling 565545, clone D8S9Y, 1:2000), rabbit anti-Lamin B1 polyclonal IgG (Proteintech 12987- 1-AP, 1:2000) and mouse anti-GAPDH monoclonal IgG (Origene TA802519, clone OTI2D9, 1:2000). Secondary antibodies were donkey anti-rabbit IgG (Alexa Fluor 790, Jackson ImmunoResearch 711-655-152) or donkey anti-mouse IgG (Alexa Fluor 680, Jackson ImmunoResearch 715-625-150), all a 1:10,000 dilution from a 1.5 mg/mL stock. Blots were quantified using Image J Software^10^.

### Cytokine Protein Analysis

Supernatants from HaCaT cells were collected, centrifuged at 4°C for 5 min at 5000 g and middle layers collected and stored at −80°C until analysis using Human CCL26/Eotaxin-3 DuoSet ELISA from R&D Systems/Bio-Techne (Cat. No. DY346, Minneapolis, MN, USA).

### Publicly Available RNA sequencing Data Analysis

Gene Expression Omnibus (GEO) data accessed for this study from publicly available datasets are as follows: GSE121212, GSE147424, and GSE153760. GSE121212 provided bulk RNA sequencing from skin biopsies, which was plotted as relative expression between control and lesional skin samples for AHR, CYP1A1, CYP1A2, CYP1B1, NFE2L2 and NQO1. GSE147424 provided scRNAseq gene expression, and the largest keratinocyte population, KC1, was used to plot normalized mean expression of AHR, CYP1A1, and CYP1B1. GSE153760 provided scRNAseq gene expression for keratinocytes in skin biopsies as well as suction blister biopsies. Single-cell RNA sequencing data were filtered to include only high-quality cells. Normalization, scaling and dimensionality reduction was performed using Seurat R package^11^, whereas integration was run with Harmony^12^. Integrated datasets were clustered with resolution 0.5. Keratinocytes were identified based on the marker genes defined in the manuscript^13^. Expression of AHR, CYP1A1 and CYP1B1 were plotted between control and lesional keratinocytes. TF binding motif enrichment analyses of identified differentially expressed genes from GSE147424 were evaluated for binding motifs using the HOMER software package^14^. Briefly, HOMER uses a library of >7,000 TF binding models (in the form of position weight matrices) to scan a set of input sequencing for statistical enrichment of each position weight matrix. Calculations were performed using ZOOPS (zero or one occurrence per sequence) scoring coupled with hypergeometric enrichment analysis to determine motif enrichment. Input enhancer sequences were also assessed for statistical enrichment of motifs for AHR binding sites using the findPeaks program and factor mode within HOMER. Significantly enriched TF binding site motifs are expressed as log p values.

### RNA Sequencing and Analysis

RNA-seq libraries were generated as previously described^15, 16^ using 500 ng of purified RNA using the Zymo Research Direct-zol RNA microprep kit (Cat. No. R2062, Irvine, CA, USA). In brief, mRNAs were enriched by incubation with Oligo d(T) Magnetic Beads (New England Biolabs, Cat. No. S1419S, Ipswich, MA, USA) and then fragmented/eluted by incubation at 94°C for 9 min. Poly A enriched mRNA was fragmented, in 2x Superscript III first-strand buffer (Invitrogen/Thermo Fisher, Cat. No. 12574026, Waltham, MA, USA) with 10 mM DTT (Thermo Scientific, Cat. No. R0861, Waltham, MA, USA), by incubation at 94°C for 9 min, then immediately chilled on ice before the next step. The 10 µL of fragmented mRNA, 0.5 µL of random primer (Invitrogen/Thermo Fisher, Cat. No. 48190011, Waltham, MA, USA), 0.5 µL of Oligo dT primer (Invitrogen/Thermo Fisher, Cat. No. 18418012, Waltham, MA, USA), 0.5 µL of SUPERase-In (Ambion/Thermo Fisher, Cat. No. AM2694, Waltham, MA, USA), 1 µL of dNTPs (10 mM, Thermo Scientific, Cat. No. R0194, Waltham, MA, USA) and 1 µL of DTT were heated at 50°C for 3 minutes. At the end of incubation, 5.8 µL of water, 1 µL of DTT (100 mM), 0.1 µL Actinomycin D (2 mg/mL, Invitrogen/Thermo Fisher, Cat. No. A7592, Waltham, MA, USA), 0.2 µL of 1% Tween-20 (Sigma-Aldrich, Cat. No. P1379, St. Louis, MO, USA) and 0.2 µL of Superscript III (Invitrogen/Thermo Fisher, Cat. No. 18080093, Waltham, MA, USA) were added and incubated in a PCR machine using the following conditions: 25°C for 10 min, 50°C for 50 min, and a 4°C hold. The product was then purified with RNAClean XP beads (Beckman Coulter, Cat. No. A63987, Indianapolis, IN, USA) according to manufacturer’s instruction and eluted with 10 µL nuclease-free water. The RNA/cDNA double-stranded hybrid was then added to 1.5 µL of Blue Buffer (Enzymatics, Cat. No. B0110, Beverly, MA, USA), 1.1 µL of dUTP mix (10mM dATP, dCTP, dGTP and 20 mM dUTP, Enzymatics, Cat. No. N2050-10-L, Beverly, MA, USA), 0.2 µL of RNase H (5 U/mL, Enzymatics, Cat. No. Y9220L, Beverly, MA, USA), 1.05 µL of water, 1 µL of DNA polymerase I (Enzymatics, Cat. No. P7050L, Beverly, MA, USA) and 0.15 µL of 1% Tween-20. The mixture was incubated at 16°C for 1 h. The resulting dUTP-marked dsDNA was purified using 28 µL of Sera-Mag Speedbeads (Cytiva, Cat. No. 65152105050250, Marlborough, MA, USA), diluted with 20% PEG8000, 2.5M NaCl to final of 13% PEG, eluted with 40 µL EB buffer (10 mM Tris-Cl, pH 8.5) and frozen at - 80°C. The purified dsDNA (40 µL) underwent end repair by blunting, A-tailing and adaptor ligation as previously described^14^ using indexed barcoding adapters (Perkin Elmer, NEXFLEX Unique Dual Indexing Barcodes). Libraries were PCR-amplified for 9-14 cycles, purified by cleanup with Speedbeads, and quantified using a Qubit dsDNA HS Assay Kit (Thermo Fisher Scientific, Cat. No. Q32854, Waltham, MA, USA). RNA-seq libraries were sequenced using PE150 and an SP flow cell on a NOVASeq 6000 at the CCHMC DNA Sequencing and Genotyping Core Facility. Sequencing data were mapped with STARR^17^ to the GRCh38 reference genome and sequencing counts were generated using HOMER analyzeRepeats.pl^14^. Differentially expressed genes were identified with DeSeq2^18^ using the HOMER wrapper script, getDiffExpression.pl. We filtered genes to those with a minimum transcript per million (TPM) value >8 (to remove low-transcript level genes), absolute fold change >1.5, and adjusted p-value <0.05 (when comparing any 2 of the 3 groups), and performed hierarchical clustering with Morpheus (Broad Institute) using a ‘one minus Pearson correlation’ and ‘average linkage’ method. Gene ontology enrichment analysis, which uses statistical methods to determine functional pathways and cellular processes associated with a given set of genes, was performed with the ToppGene suite (Cincinnati Children’s Hospital Medical Center, https://toppgene.cchmc.org/navigation/termsofuse.jsp)19. Principal Component Analysis (PCA) from gene expression data was performed using GraphPad Prism 9 (GraphPad Software, San Diego, CA, USA) with standardize method, and principal components were selected based on parallel analysis with eigenvalues greater than those from 1000 simluations at the 95^th^ percentile with auto random seed.

### Statistical Analyses

Statistical analysis of publicly available gene expression data, qRT-PCR gene expression of HaCaT cells, and cytokine protein quantification with ELISA was performed and graphed using GraphPad Prism 9 (GraphPad Software, San Diego, CA, USA).

## Results

### Disruption of AHR signaling in AD patients

We analyzed the expression of *AHR* and well-known AHR target genes such as *CYP1A1*, from a large study among publicly available data sets utilizing biopsies from AD subjects (GSE121212); Bulk RNA sequencing data of skin biopsies from 21 AD subjects (lesional skin) and 38 control subjects (healthy skin) (**Figure 1**). *AHR* expression trended toward an increase in lesional skin compared to control skin (p=0.051). In contrast, the AHR target genes *CYP1A1*, *CYP1A2*, and *NFE2L2* (*NRF2*) were all significantly decreased in lesional skin (p=1.01*10^-3^, p<1.00*10^-15^ and p=6.10*10^-8^, respectively). There was no difference in *CYP1B1* or *NQO1* when comparing control to lesional skin (p=0.096 and 0.162, respectively). These data suggest that lesional skin has decreased activation of AHR target genes, despite a trend toward increased AHR expression, and suggest that the AHR signaling pathway is dysregulated in AD skin.

**Figure 1.**
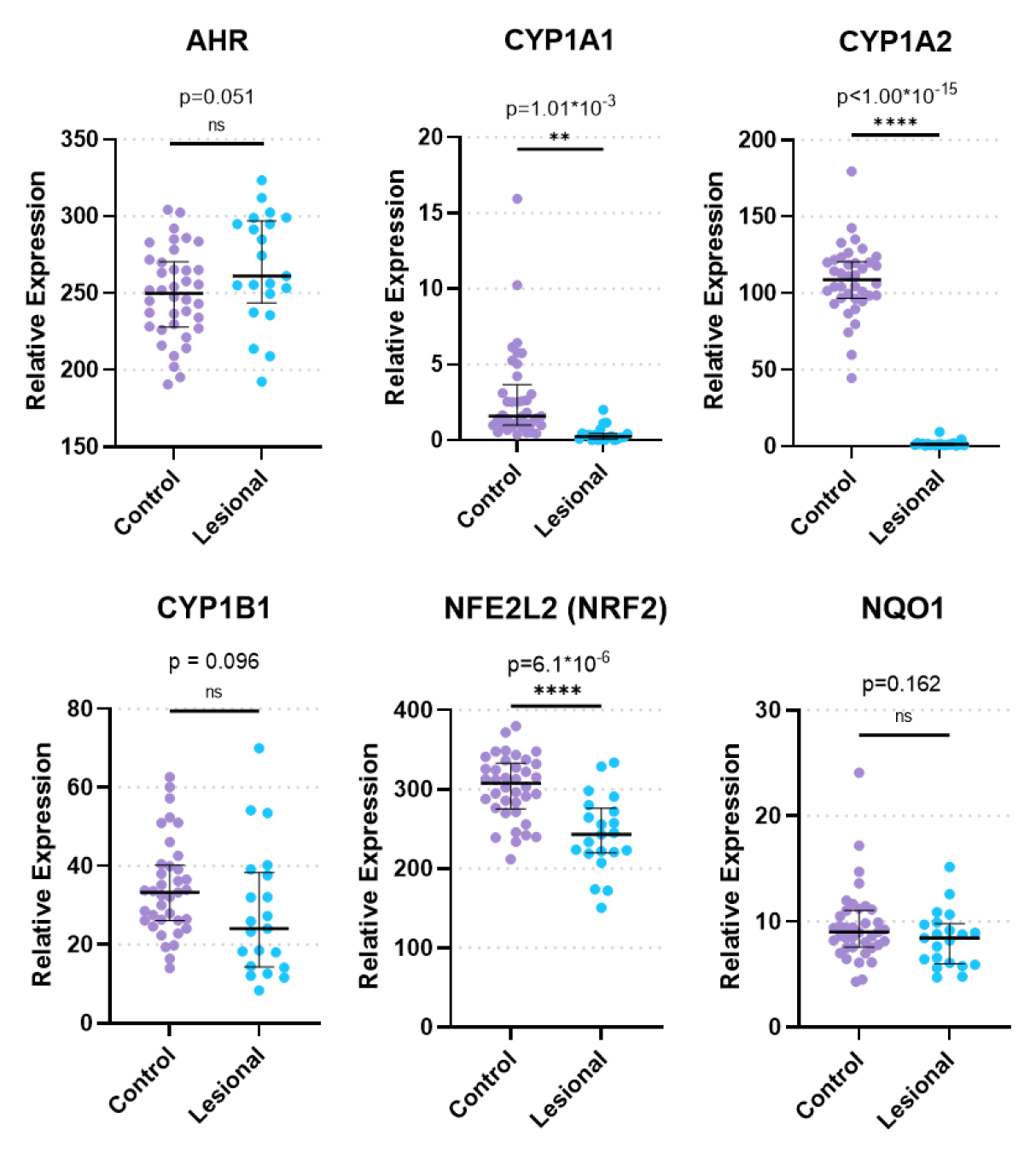
Relative Expression of AHR and Key AHR Target Genes from Bulk RNA Sequencing of Control and Lesional AD Skin Biopsies. mRNA expression of the indicated genes from GSE121212, 38 control subjects, 21 with AD. Unpaired t-test (two-tailed, α=0.05). ** = p<0.01, **** = p<0.0001.

### Altered AHR signaling in keratinocytes of subjects with AD

Focusing on keratinocytes, we analyzed publicly available single-cell RNA sequencing (scRNA-seq) data from biopsies of subjects with AD compared to healthy controls (**Figure 2**). We analyzed the largest keratinocyte population (designated KC1 in He et al. 2020^20^; GSE147424) of a scRNA-seq data from 5 AD subjects (4 lesional skin biopsies, 5 non-lesional skin biopsies) and 6 healthy control subjects (healthy skin biopsies). Lesional keratinocytes had significantly increased *AHR* expression compared to healthy control (p<0.0001) and non-lesional keratinocytes (p<0.0001). In contrast, both lesional and non-lesional keratinocytes had significantly decreased *CYP1A1* expression compared to healthy controls (p<0.001 and p<0.01, respectively). *CYP1B1* expression was decreased in non-lesional keratinocytes compared to control keratinocytes (p<0.0001) (**Fig. 2A**). We then validated these findings using a second scRNA-seq data set (GSE153760) from 8 AD subjects (4 lesional skin suction blisters, 4 lesional skin biopsies) and 7 healthy control subjects (5 healthy skin suction blisters, 2 healthy skin biopsies). Data from keratinocytes isolated from suction blisters and full biopsies were pooled within groups (AD or healthy control), and normalized expression of each keratinocyte were analyzed. *AHR* expression was increased in lesional keratinocytes compared to control (p<0.0001), though *CYP1A1* expression was not significantly different (p=0.17). *CYP1B1* expression was significantly increased in lesional keratinocytes compared to controls (p=0.017) (**Fig. 2B**).

**Figure 2.**
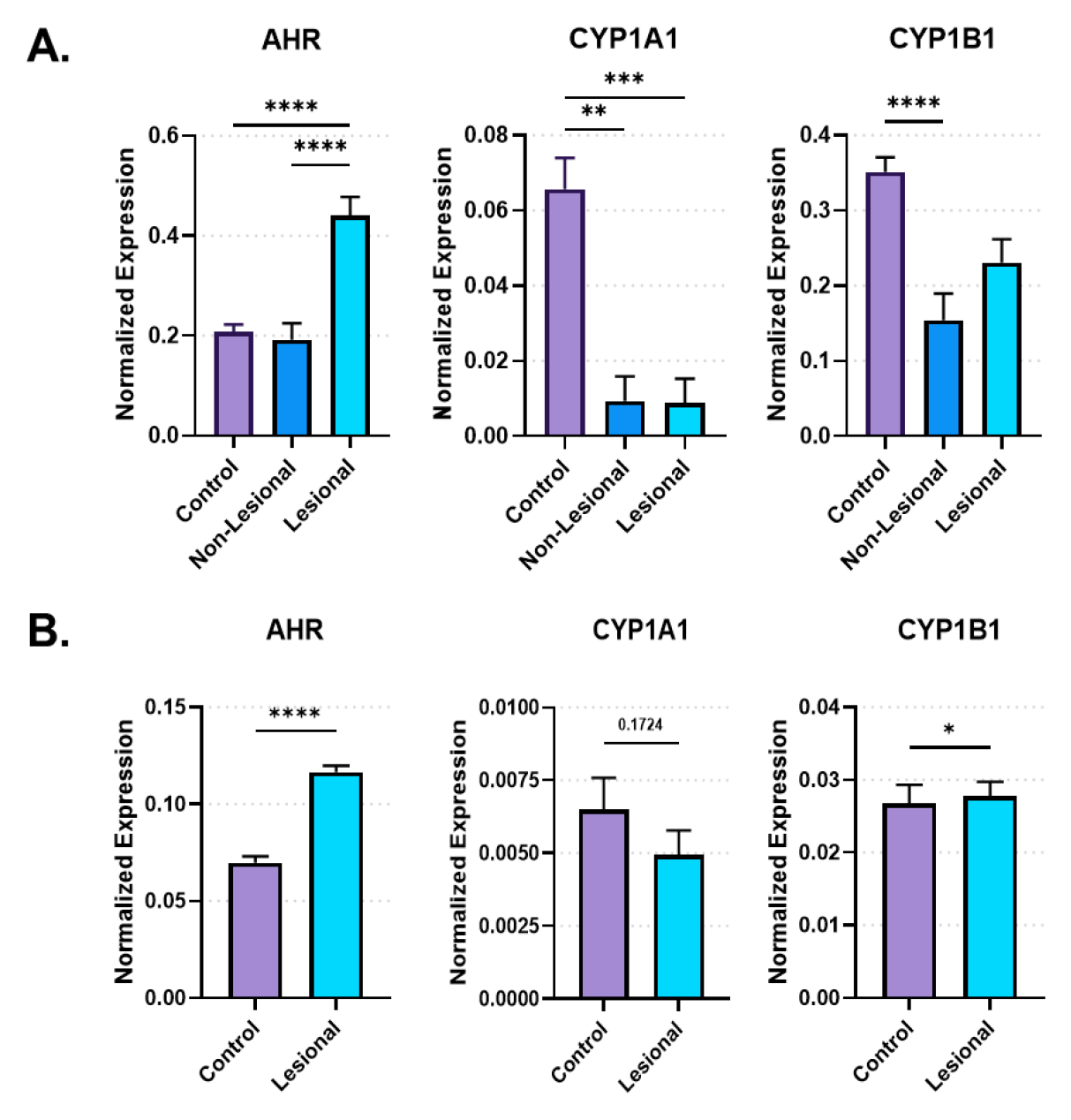
Keratinocyte-Specific Expression of AHR and Key Target Genes from Single Cell RNA Sequencing. Individual normalized KC cell expression of genes is shown; bar represents mean (+/-) standard error of the mean. **A.** From GSE147424, *AHR*, *CYP1A1*, and *CYP1B1* expression in KC1 (largest keratinocyte population) from healthy control, non-lesional AD, and lesional AD biopsies is shown. Kruskal Wallis test with Dunn’s test for multiple comparisons was used. ** = p<0.01, *** = p<0.001, **** = p<0.0001. **B.** From GSE153760, normalized mean expression of *AHR*, *CYP1A1*, and *CYP1B1* in keratinocytes from healthy control and lesional AD biopsies or suction blister samples (suction blisters pooled with biopsies from same groups) is shown. Mann-Whitney test (two-tailed, α=0.05). * = p<0.05, **** = p<0.0001

### Overlap between the gene signature of AHR-activated keratinocytes and the AD transcriptome

To investigate whether an overlap exists between AHR target genes and genes regulated in AD keratinocytes, we compared differentially expressed genes (DEGs) from keratinocytes of patients with AD (Lesional AD vs healthy controls from GSE 147424, referred here as “AD transcriptome”) with DEGs from HaCaT cells (a human keratinocyte immortalized cell line^21^) treated with an endogenous AHR ligand, FICZ for 24 hours (FICZ vs control, referred to here as “AHR transcriptome”) (**Figure 3**). A total of 290 genes were differentially expressed in the AD transcriptome, compared to 730 genes in the AHR transcriptome. Thirty seven genes overlapped between the AHR transcriptome and the AD transcriptome (12.8% of AD transcriptome; **Fig. 3A**). When the log_2_(fold change) of each of these 37 shared genes is compared, 24 of the 37 overlapped genes (approximately 2/3) were changing the direction of expression between the AD and AHR activation condition (**Fig. 3B**). Gene ontology analysis of the 37 shared genes revealed significant enrichment of biologic processes such as keratinocyte differentiation, skin development and cornification, in addition to cellular components of keratin filament, intermediate filament and connexin complex (**Fig. 3C**).

**Figure 3.**
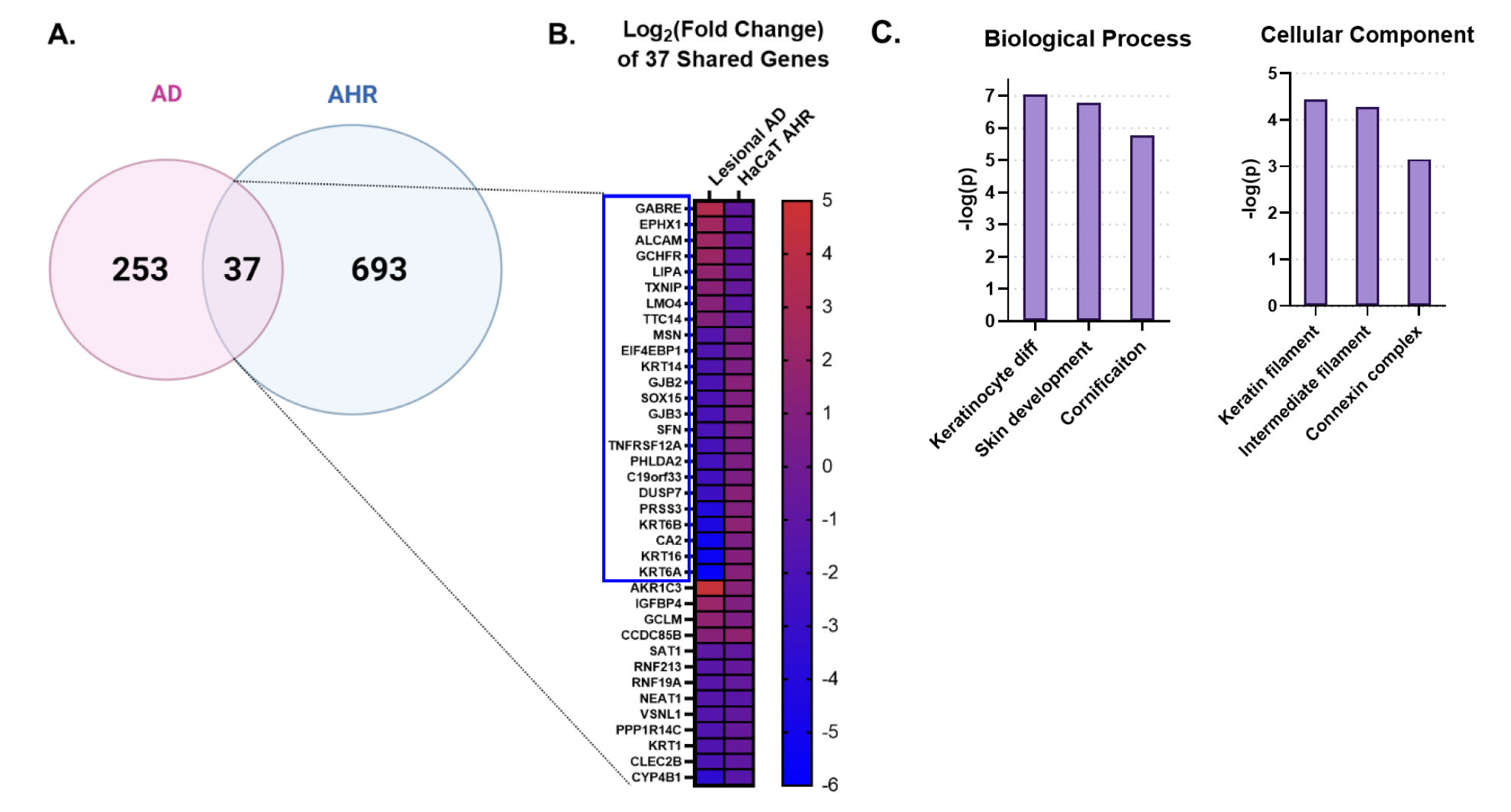
Intersecting the AD Transcriptome and AHR Transcriptome. We compared the differentially expressed genes (DEGs) from AD lesional keratinocytes (Lesional AD vs control from GSE147424) to HaCaT AHR-regulated genes (FICZ treatment vs control from HaCaT RNA sequencing). **A.** Venn diagram showing unique DEGs from keratinocytes of AD lesional biopsies (253 genes), unique DEGs regulated by AHR (693 genes), and DEGs shared by these two groups (37 genes, 12.8% of AD transcriptome). **B.** Heat map of the log2(fold change) of the 37 shared genes in (A), sorted first by genes that changed direction (blue box, 24/37 genes, ∼2/3 of genes), followed by fold change (high to low) in the AD group. **C.** Gene ontology relationships of the 37 shared genes were analyzed.

Next, we asked if AHR activation can attenuate the IL-13-mediated response. We stimulated HaCaT cells with IL-13, with and without AHR activation via FICZ. RNAseq analysis demonstrated that IL-13-regulated genes in HaCaTs (IL-13 vs Control) could be altered by addition of AHR-activation (IL-13 vs IL-13+FICZ; **Figure 4**). A total of 36 of the 344 IL-13-regulated genes (10.5% of IL-13 transcriptome) were changed by AHR activation (**Fig. 4A**). When these 36 shared genes were mapped according to log_2_(fold change), 24/36 genes (2/3 or 67%) changed direction of transcription between IL-13 and IL-13+AHR activation conditions (**Fig. 4B**). Gene ontology of these 36 shared genes revealed enrichment of extracellular matrix genes as well as inflammatory processes such as chemotaxis, humoral response, alternative complement, and IL-17 signaling (**Fig. 4C**).

**Figure 4.**
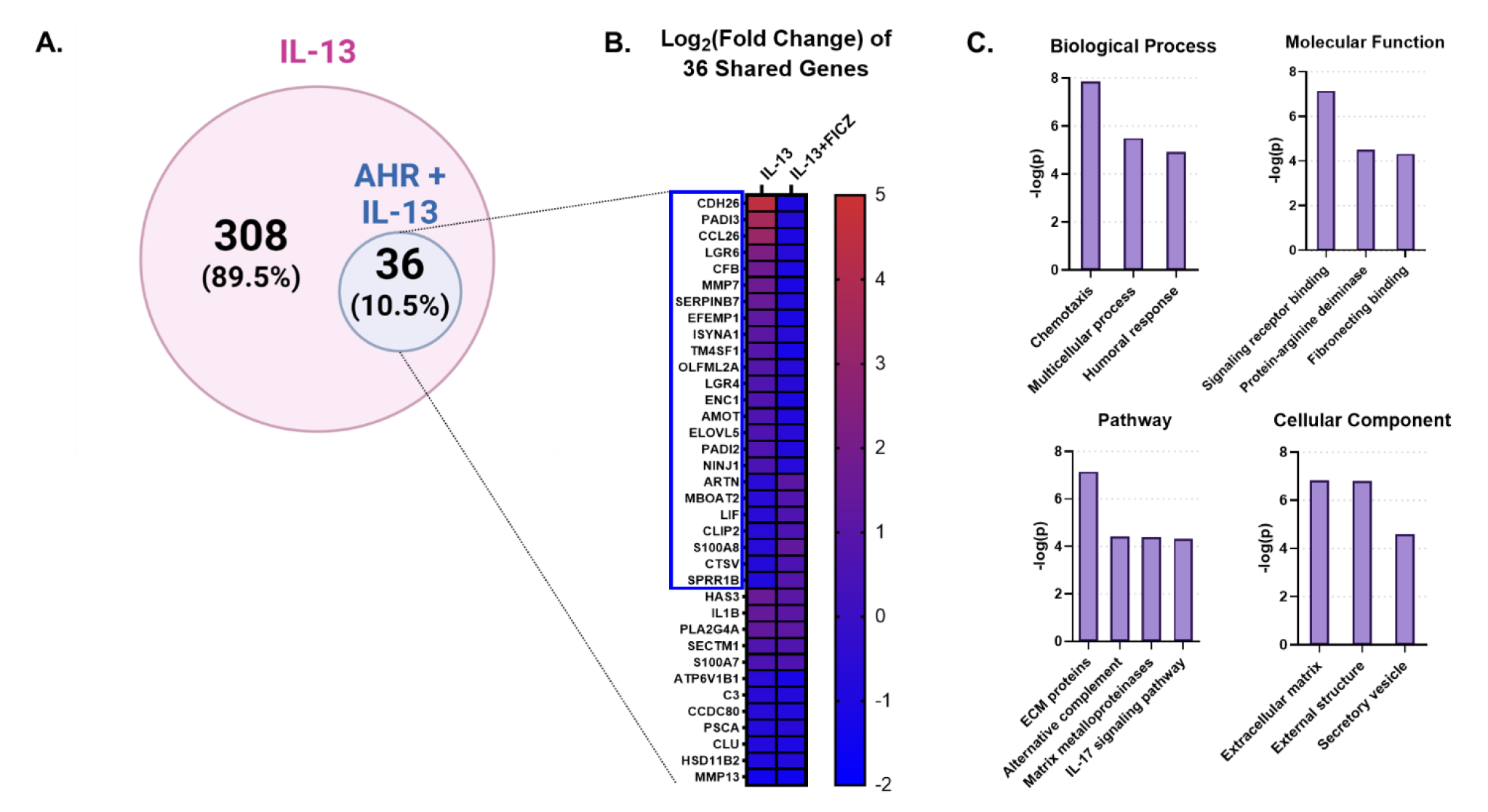
IL-13-Regulated Genes Also Regulated by AHR in HaCaT Cells. Using RNA sequencing analysis of HaCaT cells, we compared the differentially expressed genes (DEGs) from IL-13-treatment (CTL vs IL-13 [10 ng/mL] for 24 hours) to DEGs from IL-13+FICZ treatment (IL-13 [10 ng/mL] vs IL-13 [10 ng/mL] + FICZ [1 µM]) to model AD vs AD+AHR activation. **A.** Diagram showing differentially expressed genes regulated by IL-13 treatment alone (CTL vs IL-13, 308 genes) as well as a subgroup of these genes which were also regulated by AHR (IL-13 vs IL-13+FICZ, 36 genes, 10.5% of IL-13-regulated genes). **B.** Heat map of the log2-transformed fold change of the 36 shared genes in (A), sorted first by genes that changed direction (blue box, 24/36 genes, 2/3 of genes), followed by fold change (high to low) in the IL-13 group.). **C.** Gene Ontology relationships of the 36 shared genes were analyzed.

We then broadened our analysis and compared gene expression of control (untreated) HaCaTs, IL-13 treated HaCaTs and IL-13+FICZ treated HaCaTs, including only differentially expressed genes in any 2 of the 3 comparisons (CTL vs IL-13, CTL vs IL-13+FICZ, or IL-13 vs IL-13+FICZ; with adjusted p-value <0.05). Initial hierarchical clustering revealed three distinctive patterns of expression from a total of 100 genes: 1) Genes which were upregulated by IL-13 and downregulated by the addition of FICZ (IL- 13+FICZ group) (**Supplemental Fig. 1A**); 2) Low-expression genes which were downregulated by IL-13 and upregulated by the addition of FICZ (IL-13+FICZ group) (**Supplemental Fig. 1B**); and 3) High-expression genes which were downregulated by IL-13 and upregulated by the addition of FICZ (IL-13+FICZ group) (**Supplemental Figure 1C**). Principal component analysis (PCA) confirmed clustering of each treatment group from first two principal components (**Supplemental Figure 1D**). Most notable was *CCL26* (Eotaxin-3), a chemokine responsible for eosinophil chemotaxis, which is associated with AD severity and disease activity^22^, extrinsic AD in a Danish cohort^23^, and early-onset AD in children^24^. *CCL26* was one of the most upregulated genes in the IL-13 transcriptome and was reversed by FICZ. Other genes associated with allergic inflammation include *CDH26*, encoding for a cadherin protein that is increased during allergic inflammation in gastrointestinal mucosa^25^, and *FLG*, associated with the cornified envelope and barrier function. *FLG* mutations are associated with AD^26^. We further analyzed the gene ontology of these 100 genes (**Supplemental Figure 2**). The most significantly related biologic process was “skin development”, followed by epidermal-related processes, including “plasma membrane organization”, “peptide cross-linking”, “Epidermal Growth Factor Receptor (EGFR) activity”, and “epidermal cell differentiation” (**Supplemental Fig. 2A**). The most significant molecular function category was “growth factor activity”, with several other molecular functions related to signaling and fatty acid receptor binding (**Supplemental Fig. 2B**). The most significant pathway identified was “extracellular matrix (ECM)-associated proteins” and notable was inclusion of “cornified envelope formation” and “phototherapy-induced nuclear factor erythroid 2-related factor 2(NRF2)” (**Supplemental Fig. 2C**). The only two cellular components were “cornified envelope” and “AP-1 complex” (**Supplemental Fig. 2D**).

**Supplemental Figure 1.**
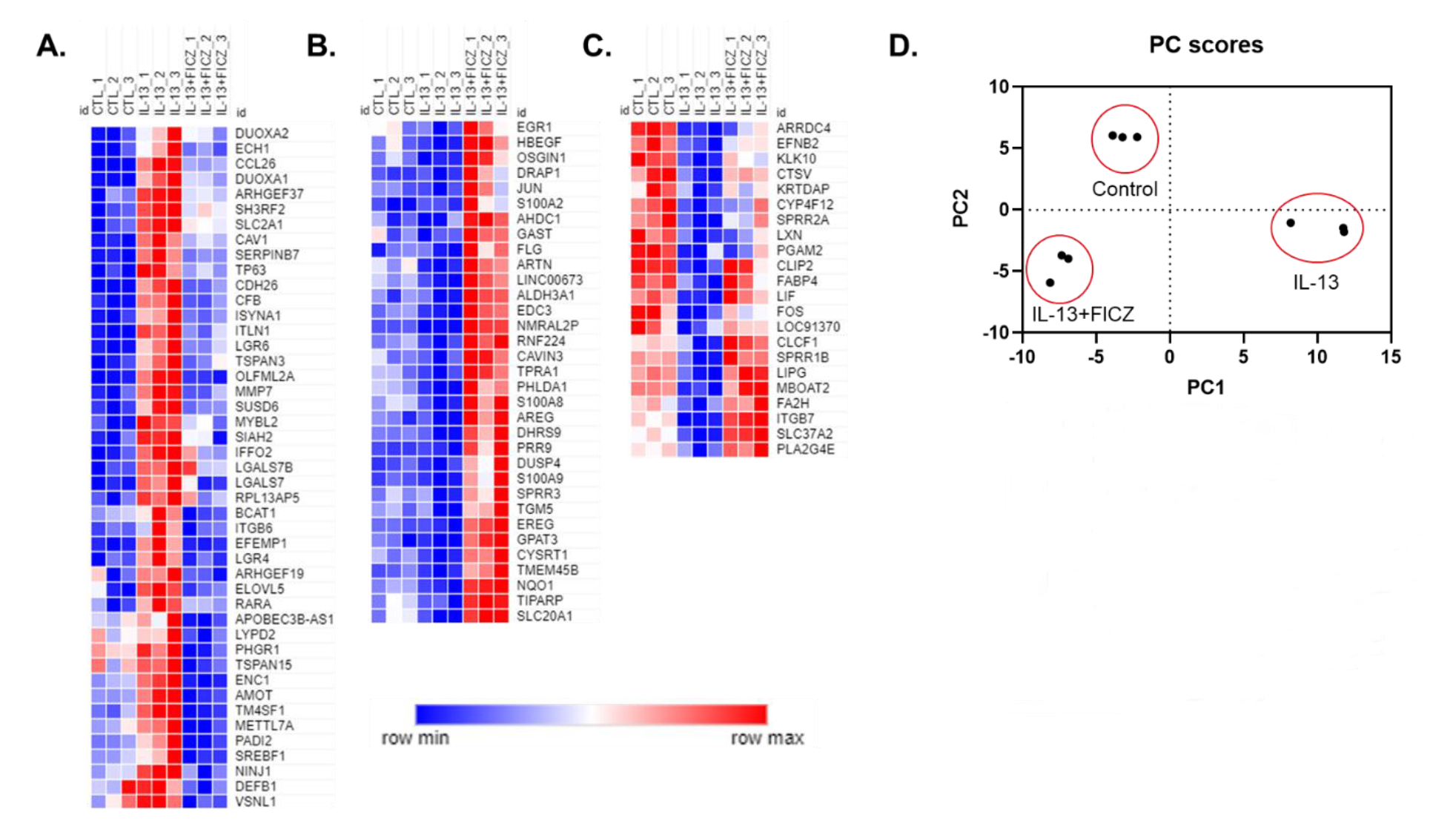
Expression patterns of differentially expressed genes in HaCaT cells treated with IL-143. and FICZ. RNA sequencing was performed using confluent HaCaT cells exposed to IL-13 (10 ng/mL) with and without FICZ (1 uM) for 24 hours. Genes were limited to those with transcript per million (TPM) >8, and only genes whose absolute fold change was >1.5 and adjusted p-value <0.05 (when comparing any 2 of the 3 groups). To better visualize patterns of expression, columns were arranged to show Control, IL-13 and IL-13+FICZ as follows: **A.** Genes which were upregulated by IL-13 and downregulated by addition of FICZ. **B.** Low expression genes which were downregulated by IL-13 and upregulated by addition of FICZ. **C.** High expression genes which were downregulated by IL-13 and upregulated by addition of FICZ. **D.** Principal Component Analysis (PCA) of gene expression showing first two principal components that clearly discern between each treatment group.

**Supplemental Figure 2.**
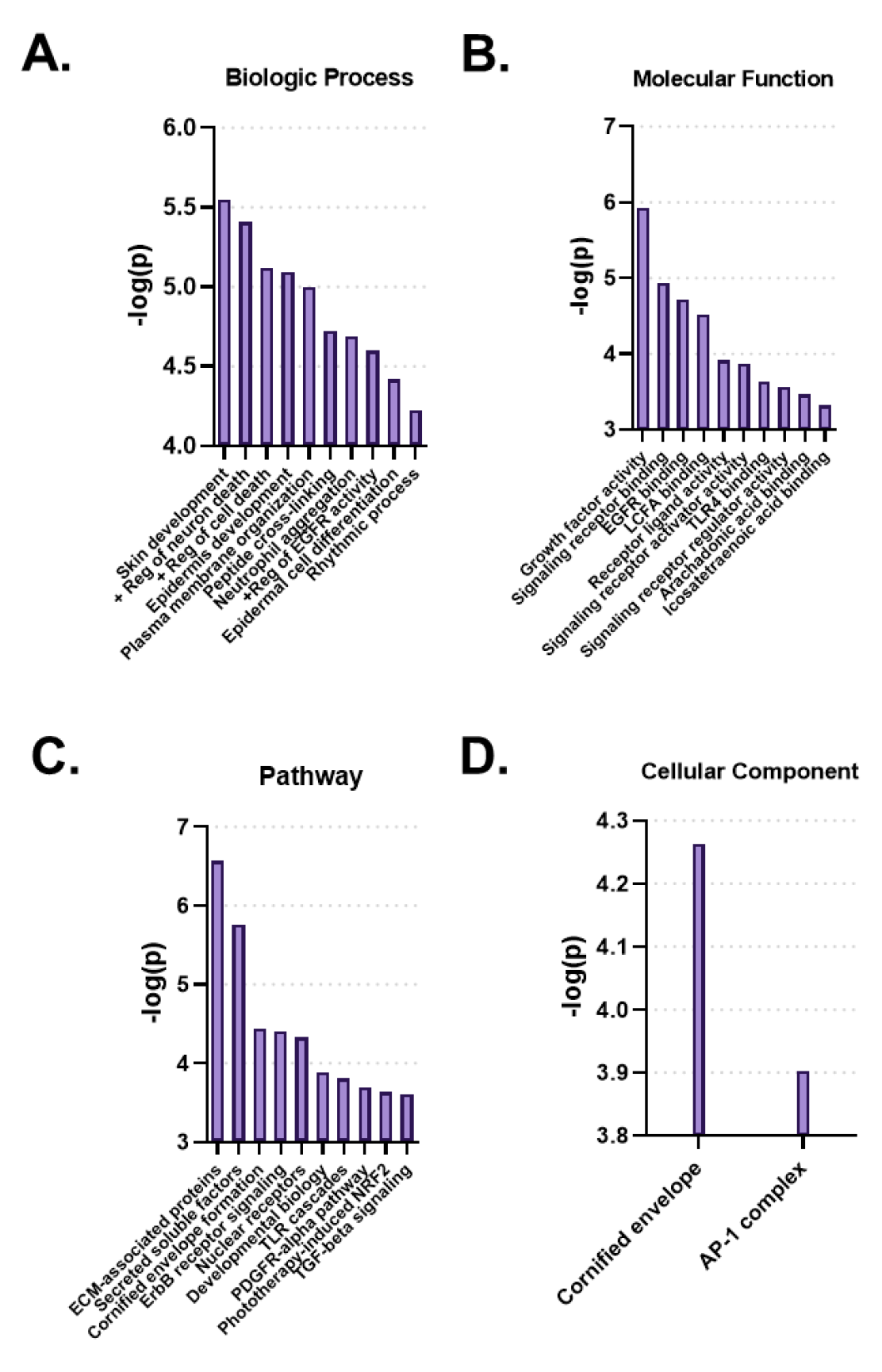
Gene Ontology Analysis of 100 DEGs Between Control, IL-13 and IL-13+FICZ Treatment. Genes that were differentially expressed between at least 2 of 3 comparisons between control, IL- 13, and IL-13+FICZ treatment (Supplemental Figure 1) were analyzed for gene ontology relationships. Shown are the top 10 biologic processes (**A**), the top 10 molecular functions (**B**), the top 10 pathways (**C**), and the top 2 cellular components (**D**). All values are expressed as -log(p-value).

### Activation of AHR signaling attenuates IL-13-mediated CCL26 expression

We performed qRT-PCR and protein analyses of CCL26 following IL-13 stimulation and activation of AHR (**Figure 5**). Analysis of canonical AHR target gene, *CYP1A1* as a positive control in HaCaT cells demonstrated that treatment with the AHR agonists FICZ and tapinarof induce *CYP1A1* expression, which was variably blunted by the AHR antagonist, GNF351 (**Fig. 5A and 5B**). When HaCaT cells were treated with IL-13, *CCL26* was induced significantly (fold change ranges from 86.4 to 1600). The induction of *CCL26* expression by IL-13 was significantly blunted by both FICZ (mean 86.4 fold with IL-13 alone vs mean 18.2 fold with IL-13+FICZ, p<0.001, **Fig. 5C**) and Tapinarof (mean 1600 fold with IL-13 alone vs mean 550 fold with IL-13 + Tapinarof, p<0.0001, **Fig. 5D**). Protein levels of CCL26 measured by ELISA under these same conditions verified the significant blunting of CCL26 protein levels by FICZ from mean of 813 pg/mL after IL-13 stimulation to 624 pg/mL after IL-13 and FICZ stimulation (p=0.029, **Fig. 5E**) and mean of 878 pg/mL following IL-13 stimulation to 663 pg/mL following IL- 13 and Tapinarof stimulation (p<0.0001, **Fig. 5F**), though the absolute amount of protein was only decreased by approximately 20%. These results validate that HaCaTs are responsive to AHR and IL-13, and that IL-13- dependent CCL26 expression and protein levels could be blunted by AHR activation.

**Figure 5.**
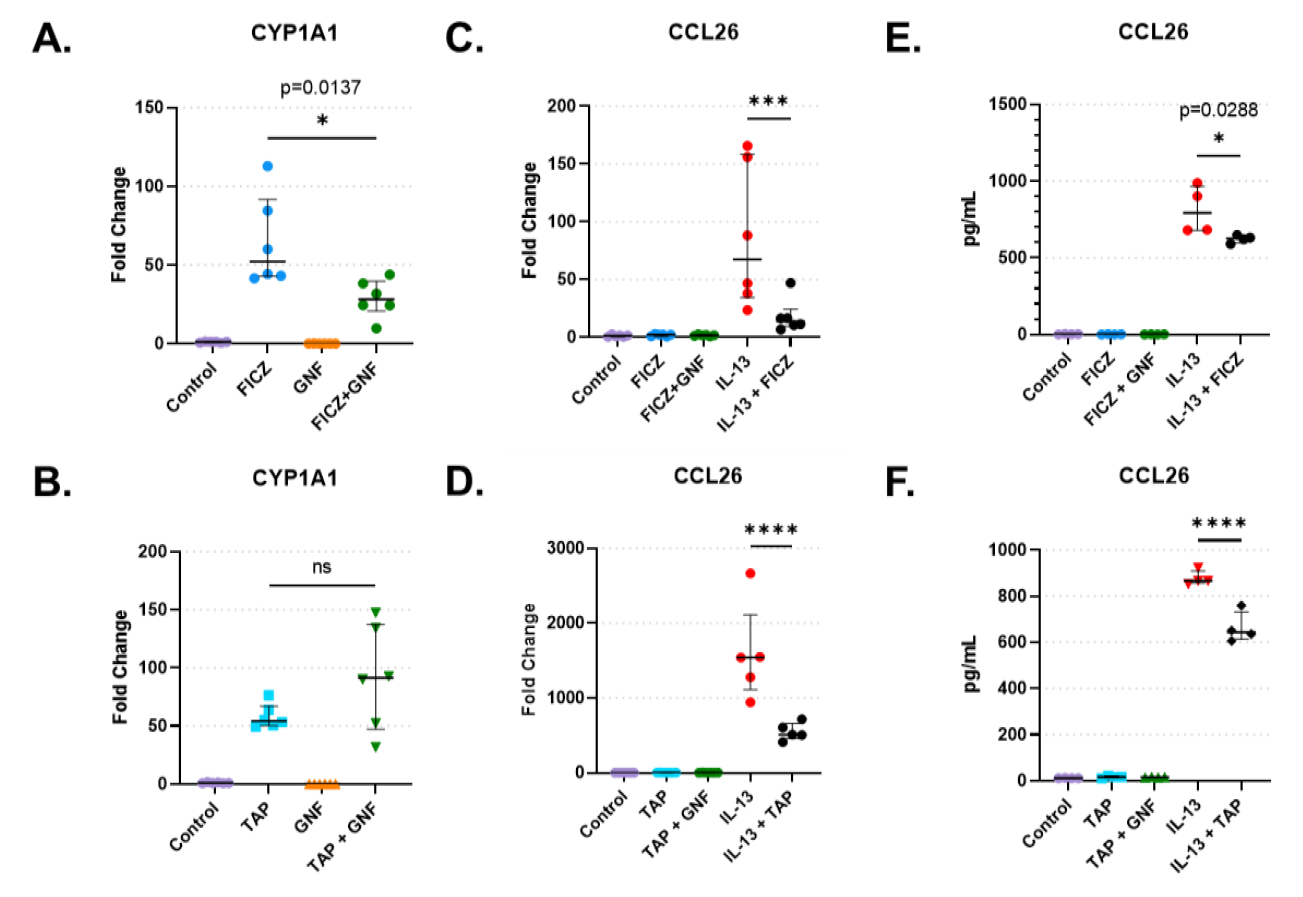
HaCaT Cell Responses to AHR activation, AHR blockade, and IL-13. **A.** CYP1A1 expression in HaCaT cells treated with AHR endogenous ligand FICZ (1 µM) (Tapinarof 1 µM in [**B.**]), and/or GNF351 (1 µM, AHR blocker) (n=6). **C.** CCL26 expression in HaCaT cells treated with FICZ (1 µM) (Tapinarof 1 µM in [**D.**]) (n=6). **E.** CCL26 protein level in HaCaT cells treated with FICZ (1 µM) (Tapinarof 1 µM in [**F.**]) (n=4). One-Way ANOVA with Tukey’s correction for multiple comparisons; * = p<0.05, ** = p<0.01, *** = p<0.001, **** = p<0.0001.

### AHR activation does not alter total STAT6, pSTAT6, or STAT6 nuclear translocation

We hypothesized that AHR mediates its effect on the IL-13 response by regulating the activity of STAT6 (a downstream transcription factor activated by IL-13 and IL-4 which regulates CCL26 expression^27^. To confirm that AHR is activated by AHR, we analyzed AHR cellular localization after 24 hours of FICZ treatment with or without GNF351. AHR was mainly present in the cytosol in both control (1% DMSO) and FICZ+GNF351-treated HaCaT cells, whereas the cytoplasmic fraction was significantly decreased (and nuclear fraction increased) in the FICZ-treated HaCaT cells (**Supplemental Fig. 3A**). Densitometry of nuclear and cytoplasmic fraction of AHR (**Supplemental Fig. 3B**) show that FICZ treatment dramatically increases nuclear translocation of AHR and that IL-13 does not affect this translocation.

**Supplemental Figure 3.**
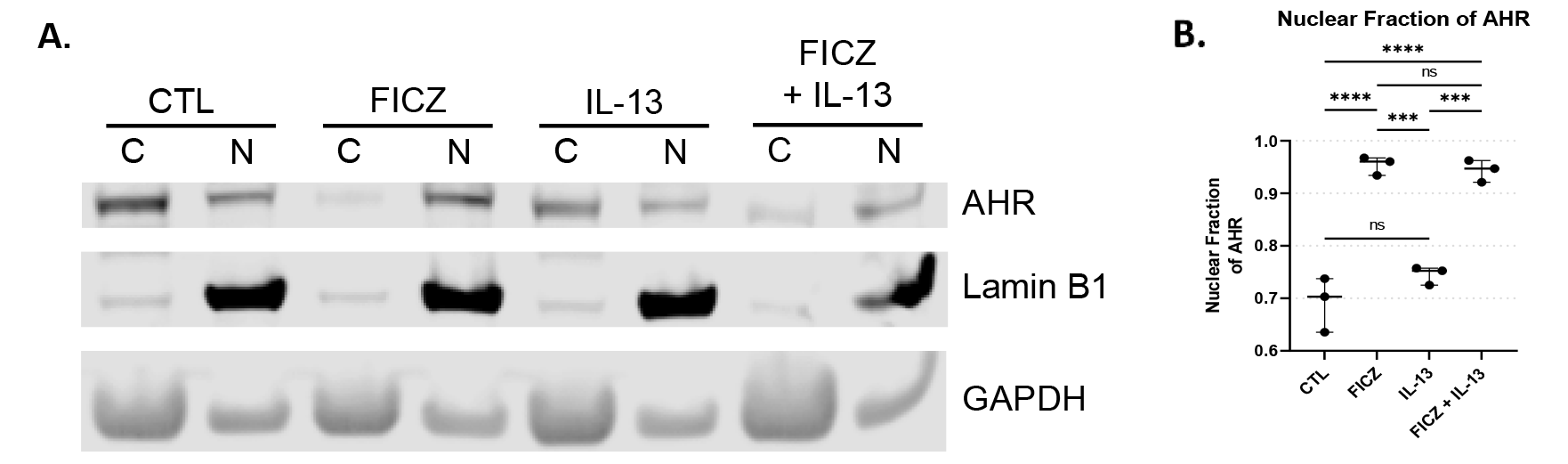
AHR Western Blot in HaCaT cells. **A.** Representative AHR western blot using cytoplasmic and nuclear fractions of the same sample after 1 hour of treatment, with nuclear marker LaminB1 and cytoplasmic marker GAPDH. Control is untreated, FICZ is 1 µM, IL-13 is 10 ng/mL, with “C” designating cytoplasmic and “N” designating nuclear fractions. **B.** Intensity of bands from A were quantified, and the normalized nuclear AHR fraction is shown (nuclear AHR / (nuclear AHR + cytoplasmic AHR)). Cytoplasmic AHR was normalized to cytoplasmic GAPDH and nuclear AHR was normalized to nuclear LaminB1. Median +/- interquartile range shown. ANOVA with Tukey’s test for multiple comparisons (n=3); *** = p<0.001, **** = p<0.0001.

To determine whether AHR activation impacts STAT6 phosphorylation or nuclear translocation, we performed western blots for pSTAT6 (**Figure 6**). pSTAT6 was not present in untreated or FICZ-treated samples and can be seen in both cytosolic and nuclear fractions of the IL-13 treatment group and the IL- 13+FICZ treatment group (**Fig. 6A**).

**Figure 6.**
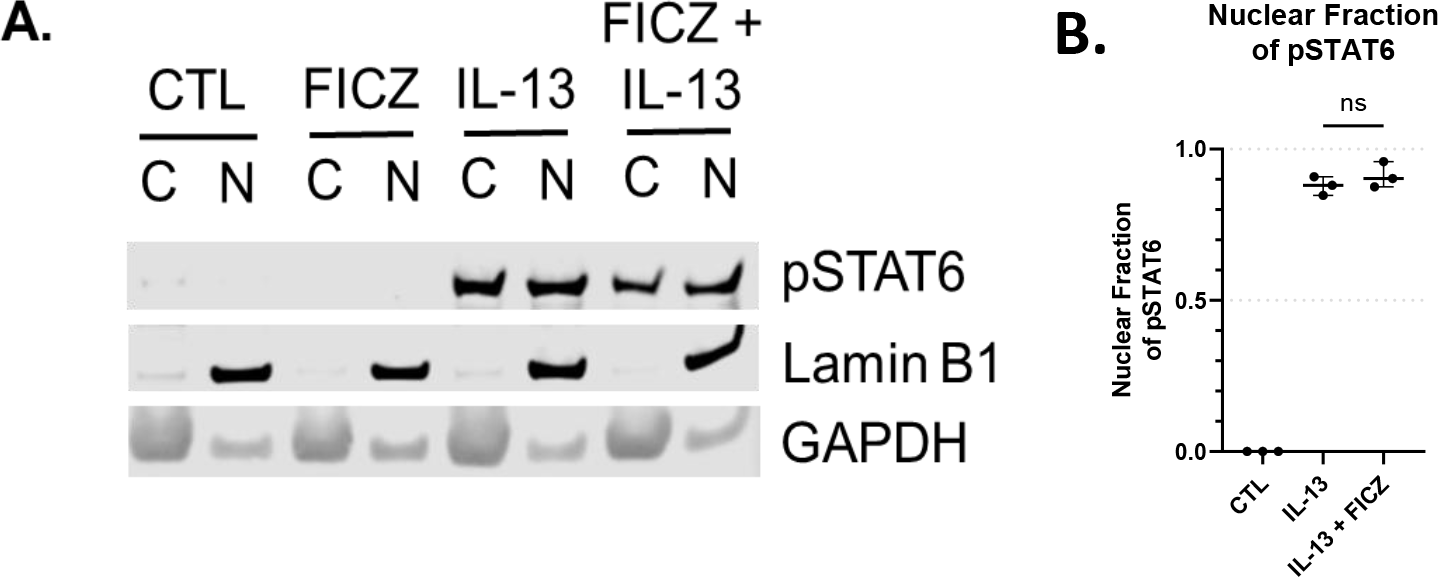
Western Blot of pSTAT6 in HaCaT cells. **A.** Representative pSTAT6 western blot using cytoplasmic and nuclear fractions of the same sample after 1 hour of treatment, with nuclear marker LaminB1 and cytoplasmic marker GAPDH. Control is untreated, IL-13 is 10 ng/mL, with “C” designating cytoplasmic and “N” designating nuclear fractions of the same sample, respectively. **B.** Intensity of bands from A were quantified, and the normalized nuclear pSTAT6 fraction is shown (nuclear pSTAT6 / (nuclear pSTAT6 + cytoplasmic pSTAT6)). Specifically, cytoplasmic pSTAT6 was normalized to cytoplasmic GAPDH and nuclear pSTAT6 was normalized to nuclear LaminB1. Median +/- interquartile range shown (n=3).

Densitometry of bands showed that the nuclear fraction of pSTAT6 was not different between IL-13 and IL-13+FICZ treated cells, indicating that AHR activation did not alter nuclear translocation of pSTAT6 (**Fig. 6B**). When total STAT6 levels were measured, only IL-13 caused an increase in total STAT6 levels, and AHR activation with FICZ did not affect total STAT6 **(Supplemental Figure 4).**

**Supplemental Figure 4.**
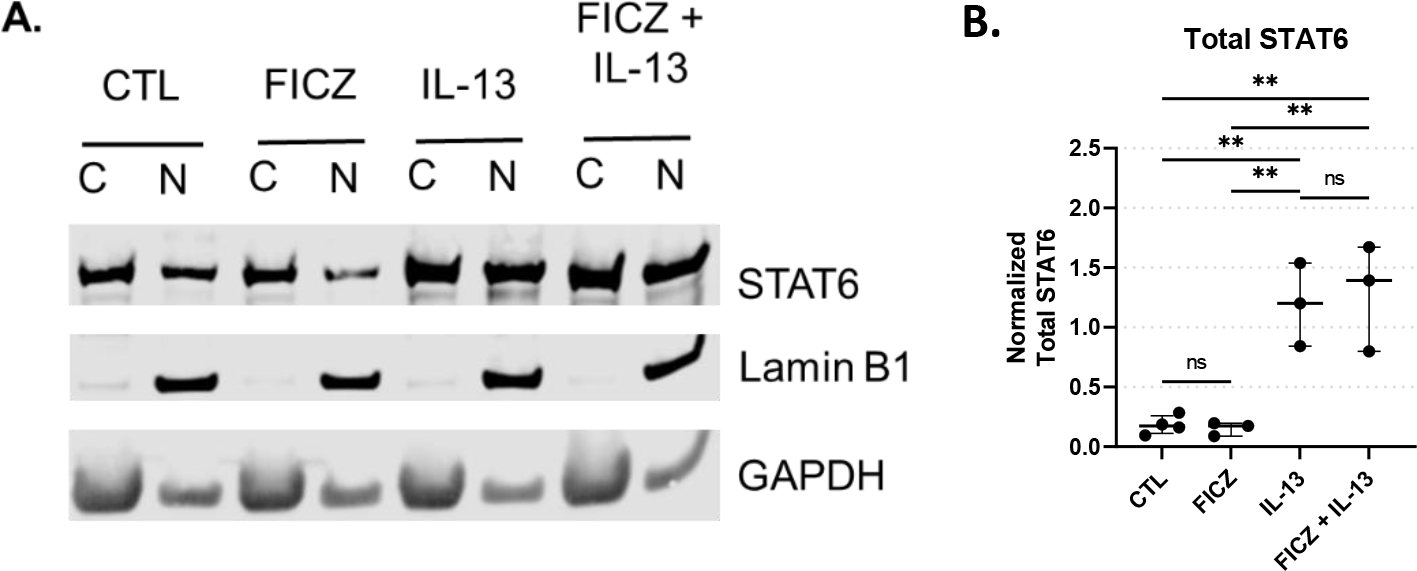
Western Blot of Total STAT6 in HaCaT cells. **A.** Representative STAT6 western blot using cytoplasmic and nuclear fractions of the same sample after 1 hour of treatment, with nuclear marker LaminB1 and cytoplasmic marker GAPDH. Control is untreated, FICZ is 1 µM, IL-13 is 10 ng/mL, with “C” designating cytoplasmic and “N” designating nuclear fractions. **B.** Intensity of bands from A were quantified, and the normalized sum of STAT6 is shown (cytoplasmic STAT6 was normalized to cytoplasmic GAPDH and nuclear STAT6 was normalized to nuclear LaminB1). Median +/- interquartile range shown (n=3). ANOVA with Tukey’s test for multiple comparisons; ** = p<0.01.

### Enrichment of AHR binding motifs in active chromatin regions of DEGs from keratinocytes of patients with AD

Because AHR activation did not affect pSTAT6 nuclear translocation, and given the overlap of differentially expressed genes between IL-13 and AHR, we tested whether STAT6 competes with AHR on DNA binding as a potential mechanism of AHR-induced changes to IL-13-regulated genes. Utilizing available scRNA-seq data from keratinocytes of patients with AD (lesional sample) compared to healthy controls (from GSE147424), we identified the most differentially expressed genes (top 20% of absolute fold change, called “DEGs”) and genes that were not differentially expressed (lowest 20% of absolute fold change as internal control genes, called “Non-DEGs”). Of these genes, we analyzed their likelihood to have active chromatin by intersecting with ChIP-seq data from human keratinocytes (GSM5113883, GSM4025776 and GSM5330873). BedTools intersect was used to determine whether any part of the gene body overlapped with areas of active chromatin as determined by H3K27 acetylation (H3K27ac) (**Figure 7**). As expected, tag density of H3K27ac marked active regions of chromatin were and centered around the transcriptional start site (TSS) of each gene (**Fig. 7B**). We found significantly more AD DEGs overlapped with H3K27ac enriched active chromatin regions compared with the Non-DEGs (94.7% vs 71.0%, p=0.038) (**Fig. 7C**). Finally, H3K27ac enriched gene regions were analyzed for consensus AHR binding motifs. DEGs contained significantly higher proportion of AHR binding motifs than Non-DEGs (39.3% vs 30.9%, p=0.007) (**Fig. 7D**). Collectively, these data suggest that AHR signaling may attenuate IL-13 induced response in keratinocytes by competing with STAT6 on DNA binding.

**Figure 7.**
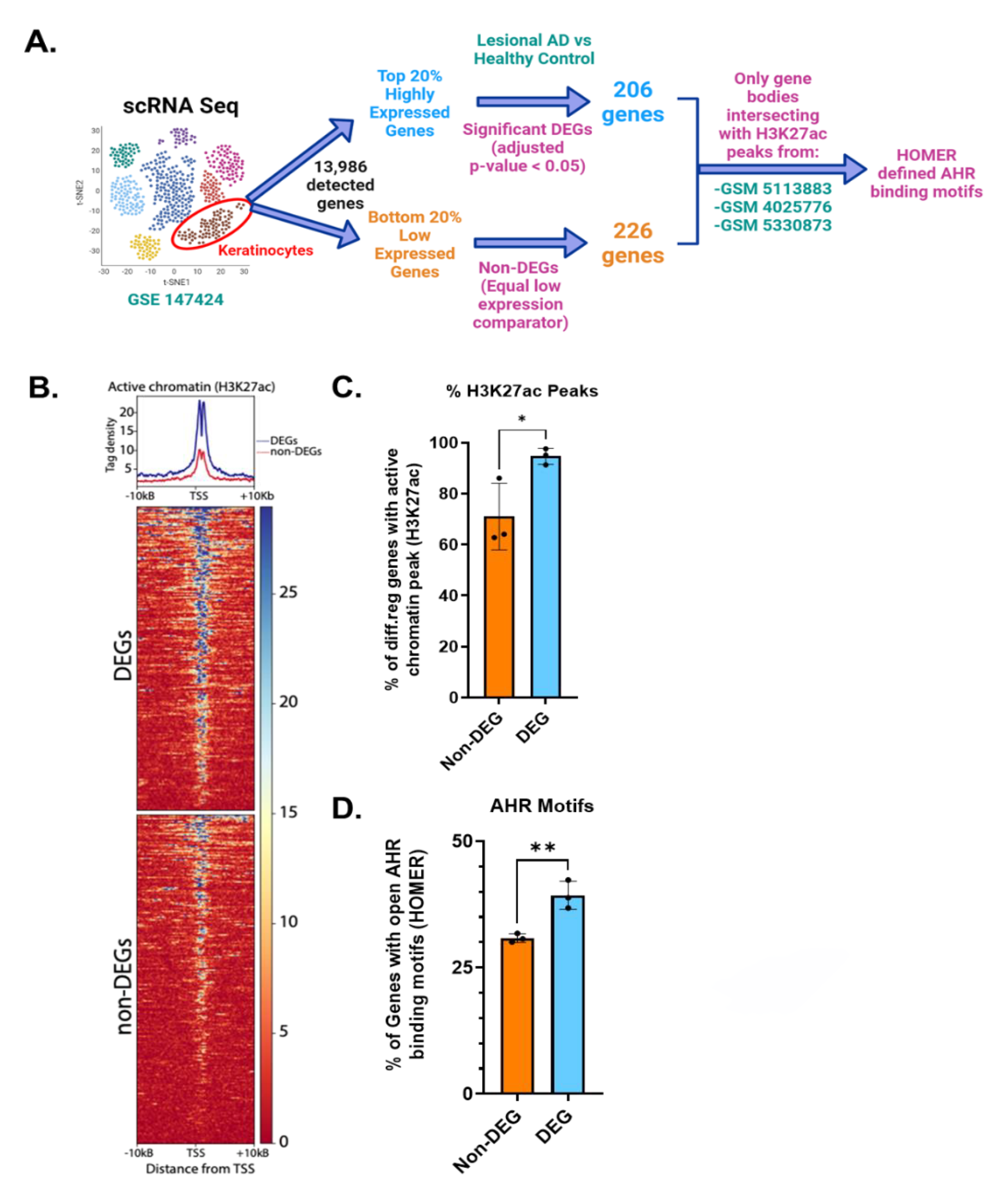
Analysis of AHR Binding Motifs in Open Chromatin Regions (H3K27ac) of Differentially Expressed Genes from Keratinocytes of Patients with AD vs Healthy’ Controls. **A.** Keratinocytes from lesional AD vs healthy controls were identified from scRNA-seq data (GSE147424, hypothetical tSNE plot shown). Next, gene lists for the Top 20% of differentially expressed genes (adjusted p-value <0.05, 206 genes) and Bottom 20% of Non-DEGs (p-value >0.05, 226 genes as internal control) were generated. **B.** H3K27ac tag density (using H3K27ac ChIP-seq peaks from 3 different human keratinocyte data sets: GSM5113883, GSM4025776 and GSM5330873) were compared in DEGs vs Non-DEGs. **C.** Percentage of DEGs that intersected with an active chromatin peak of DEGs vs Non-DEGs. **D.** Percentage of genes with AHR binding motifs between DEGs vs Non-DEGs. Unpaired t-test (n=3); * = p<0.05, ** = p<0.01.

## Discussion

In this study, we presented a collective line of evidence demonstrating that AHR signaling is altered in keratinocytes of AD patients compared to controls. We further demonstrated that activation of AHR signaling by delivery of AHR ligands induces epithelial barrier genes and attenuates IL-13-mediated induction of key AD signature genes including CCL26. Aiming to elucidate the mechanism by which AHR attenuates the IL-13 response, we hypothesized that AHR interacts and restrains STAT6. We demonstrated that AHR does not interfere with total STAT6 protein expression, STAT6 phosphorylation, or nuclear translocation, and present evidence that AHR may interfere with STAT6 at the level of DNA binding to shared target genes (summarized in **Figure 8**).

**Figure 8.**
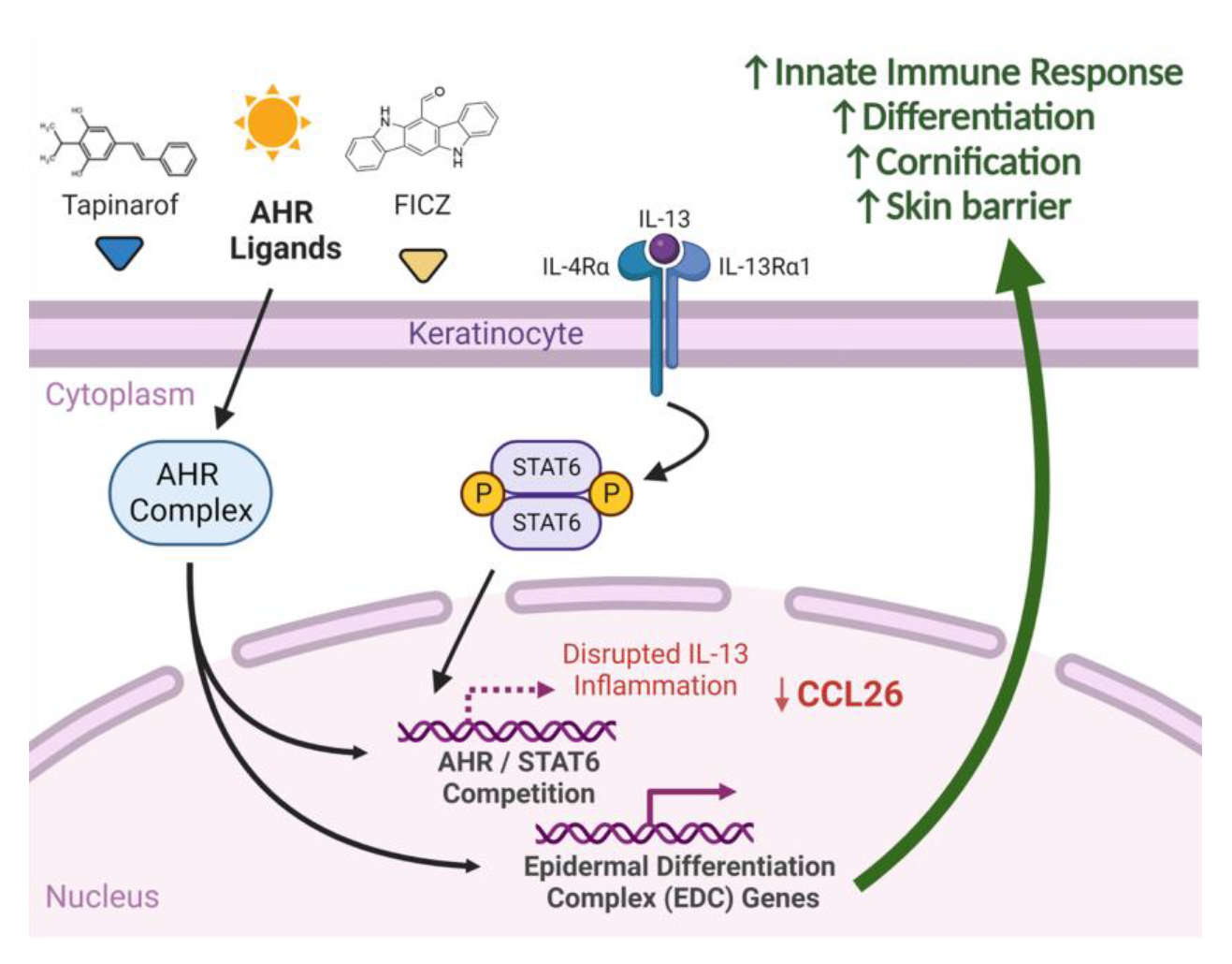
Summary of AHR and IL-13 Signaling in Keratinocytes. In keratinocytes exposed to AHR agonists and IL-13, Type 2 inflammation is disrupted and cytokines such as CCL26 are decreased, whereas genes affecting differentiation and skin barrier are upregulated, which may explain some of the efficacy of AHR agonists in atopic dermatitis.

It was unknown from prior studies whether AHR signaling was altered in keratinocytes of subjects with AD. Only recently have technologies allowed cell-type specific resolution of gene expression from biopsies to enable this analysis. Furthermore, while AHR activation was known to induce epidermal differentiation complex genes such as filaggrin and decrease inflammatory cytokines such as TSLP and CCL26, the mechanisms of these changes are not well understood.

Prior to scRNA sequencing, bulk RNA sequencing and immunohistochemistry were the best techniques available to locally probe gene expression in skin of subjects with AD. A recent study of bulk RNA sequencing of biopsies from AD patients at different ages by Renert-Yuval et al. showed decreased AHR expression in normal healthy skin, compared with lesional and non-lesional skin, in children 0-5 years old (n=17)^28^. Since the data we used for our analysis was obtained from adult patients, further study of age-associated AHR expression would be required to reconcile these differences seen between bulk RNA sequencing results. These differences also highlight the importance of single-cell resolution of gene expression afforded by newer technologies. Our analysis of keratinocyte-specific gene expression confirms the general trend seen in immunohistochemistry and bulk RNA sequencing where AHR expression is increased in lesional skin without activation of canonical AHR target genes, demonstrating that this pathway is abrogated in AD keratinocytes and may contribute to the pathology of AD^29, 30^.

Keratinocytes are uniquely positioned to have exposure to AHR ligands and precursors of AHR ligands. First, the skin is exposed to UV radiation, required for FICZ generation from endogenous tryptophan^31^. Another source of AHR ligands for keratinocytes are commensal microbes, which produce a variety of indoles and related AHR ligands in the normal course of their metabolism^32^. Notably, subjects with AD have altered microbiota (typically switching from Staphylococcus epidermitis-dominant to Staphylococcus aureus-dominant). This calls for more detailed studies of AHR ligand production by these commensal bacteria. Further, impaired barrier function of AD keratinocytes can alter exposure to environmental AHR ligands, both from commensal bacteria and also from pollutants (namely particulate matter and products of combustion). Given the importance of environmental factors on AD risk, and the ability for AHR to interact with many environmental ligands, it is plausible that AD pathogenesis relies on variable exposures to AHR ligands and precursor molecules (some protective, some pathogenic). Further study is needed to put AHR ligand exposures into context, though it is intriguing to consider AHR as a key piece of the environmental puzzle of AD and the rising incidence of other allergic diseases^33^.

Our study is the first to directly compare lesional DEGs from AD keratinocytes to AHR activation of in-vitro keratinocyte cell model (Figure 3). Shared genes are mostly reversing direction in expression between AD and AHR activation states and are involved with keratinocyte development and skin barrier function, which support this as a possible mechanism by which AHR activation might improve AD. Other studies to date support this relationship between AHR activation and improved skin barrier function. Van den Bogaard et al. used skin organoids derived from primary keratinocytes of patients with AD and showed that coal tar acted through AHR to restore filaggrin expression and other markers of keratinocyte differentiation^34^. Tsuji et al. showed that in primary human keratinocytes, AHR activation with FICZ and Glyteer (soybean tar) could induce the expression of the barrier gene filaggrin (FLG), and that FICZ and Glyteer can restore the FLG expression that was reduced by IL-4 treatment^35^. Notably, filaggrin is a key protein involved in the skin barrier and mutations in FLG are associated with AD^36^. Tapinarof has been shown by Smith et al. to induce expression of epidermal differentiation genes in keratinocytes as well as to improve inflammation in an AHR-dependent manner using both a human ex-vivo air liquid interface culture model and an imiquimod dermatitis mouse model^5^.

The crosstalk between AHR and IL-13 does not appear to be limited to just skin barrier function; we compared in-vitro AD-like conditions with and without AHR activation, and demonstrated that the shared DEGs include not only skin barrier function (extracellular matrix, external structure) but also inflammatory processes such as chemotaxis, humoral response, alternative complement, and IL-17 signaling (Figure 4). Other studies to date have also shown the ability of AHR to regulate type 2 inflammation in epithelial cells. AHR has been shown to bind to the TSLP promoter to downregulate its expression in mouse keratinocytes, and while TSLP was not among the several DEGs found in our study, the overall mechanism would be consistent with our findings for CCL26^37^. Another study revealed relevance of AHR signaling in proton pump inhibitor (PPI)-responsiveness of esophageal epithelial (EPC2) cells and reversal of approximately 20% of the IL-13 transcriptome, consistent with some of the expression reversals we describe here^38^. In another study, coal tar activation of AHR prevented IL- 4 and IL-13-dependent CCL26 expression in human primary keratinocytes derived from AD patients, consistent with our findings^34^. Of note, they showed that pSTAT6 was decreased after addition of coal tar to skin organoids pre-treated with both IL-4 and IL-13. This is in contrast to our results, where no changes to pSTAT6 were noted, though there were also differences between AHR agonist (FICZ vs coal tar) and Th2 stimulation (IL-4 and IL-13 vs IL-13 alone). Further studies will resolve more of the dynamic relationship between AHR and STAT6 signaling pathways in keratinocytes, but overall evidence is mounting that crosstalk in these two pathways is likely to play a role in AD.

It is important to recognize that AHR activation is not exclusively beneficial to skin, as studies utilizing the AHR ligands TCDD and PCBs (in addition to case studies of human exposures) demonstrate intense cystic dermatitis and inflammation (termed “chloracne” in humans)^39^. In mice, a constitutively active AHR mutant leads to a dermatitis phenotype^40^. One factor likely contributing to this differential response to AHR activation is the metabolism and stability of the many known AHR ligands; for example, TCDD has a half-life between 5-10 years in the human body^41^ compared to FICZ’s half-life of a few hours^42, 43^. Another factor complicating the study of AHR activation in the skin is the role of AHR signaling on immunologically active cells. The importance of AHR to T-helper 17 (Th17) cells is well known, and its effects on innate lymphoid cells, dendritic cells, macrophages and others cannot be discounted^44^. However, we suggest that the role of AHR in keratinocytes by endogenous ligands like FICZ is particularly important in AD. Unraveling the nuances of AHR activation as a therapeutic target in AD will require more detailed mechanistic studies controlling for the many AHR ligands, cell types, and even AD endotypes.

Our study is limited by the small number of subjects used in some of the RNA sequencing studies of keratinocytes, and this data will evolve as data from larger studies will be published. Additionally, use of keratinocytes from biopsies of subjects with AD treated with and without AHR ligands such as tapinarof would provide much more direct comparisons of the effects of AHR activation in AD than in-vitro models. We acknowledge that more complicated models such as air-liquid-interface or skin organoids may prove to be more representative of human skin, though future studies can explore this further. Finally, more in-depth studies of DNA binding competition between AHR and pSTAT6 are needed to understand crosstalk between these two transcription factors in AD.

In conclusion, we provide evidence that AHR signaling is disrupted in keratinocytes of subjects with AD, and an overlap exists between the genes that are altered by both IL-13 and genes that are regulated by AHR activation. These overlapped genes are enriched with genes with skin barrier function and innate immune response, including CCL26. AHR activation does not appear to alter STAT6 levels, activation, or nuclear translocation, and we provide evidence that DNA binding may be the primary means by which AHR activation alters IL-13 signaling. Further study of AHR in AD will help clarify to what extent this environmental sensor can regulate responses to allergic disease.

## Competing interests

The authors declare that they have no competing interests.

## Abbreviations

AD: atopic dermatitis
AHR: aryl hydrocarbon receptor
DEG: differentially expressed gene
DMSO: dimethylsulfoxide
EGFR: epidermal growth factor receptor
FICZ: 6-formyl-indolo[3,2-b]carbazole
H3K27ac: Histone 3 lysine 27 acetylation
IL: interleukin
KC: keratinocyte
NRF2: nuclear factor erythroid 2-related factor 2
PCA: principal component analysis
PCB: polychlorinated biphenyl
PPI: proton pump inhibitor
scRNA-seq: single cell RNA sequencing
STAT6: signal transducer and activator of transcription 6
TCDD: 2,3,7,8-tetrachlorodibenzo-p-dioxin
Th2: T-helper type 2

## Acknowledgements

This work is supported by the National Institute of Environmental Health Sciences (NIEHS) grant T32 ES010957 and the National Heart, Lung, and Blood Institute (NHLBI) grant R01 HL132344, and, in part, by supported, in part, by the Cincinnati Children’s Hospital Medical Center Research Foundation. Some figures for this paper were created through BioRender.com.

## Notes

### Competing Interest Statement

The authors have declared no competing interest.

